# NOTES ON THE BIOMETRY, ORIENTATION AND SPATIAL LOCALIZATION OF THE NESTS OF THE GLOBALLY NEAR-THREATENED STRAIGHT-BILLED REEDHAUNTER (*LIMNOCTITES RECTIROSTRIS*)

**DOI:** 10.1101/128553

**Authors:** Maycon S. S. Gonçalves, Priscila dos S. Pons, Christian B. Andretti, Felipe C. Bonow

## Abstract

We report the records of three nests of Straight-billed Reedhaunter, *Limnoctites rectirostris*, in southernmost Brazil. For each nest, we collected information about the biometry, spatial localization within the patch (border or inland) and cardinal orientation of the incubation chamber. Information on number of eggs and reproductive success is also presented. All nests were found early in September. The three nests were situated on the edge of patches and guided incubation chamber opening between north and east. All results are discussed. The information presented herein extends the scarce knowledge about the breeding biology of the Straight-billed Reedhaunter.

## Introduction

The Straight-billed Reedhaunter (*Limnoctites rectirostris*) is an aquatic Passeriform that inhabits humid ecosystems dominated by Eryngo *Eyngium pandanifolium* Cham. & Schltdl. (Apiaceae) of South America. This ecosystem is popularly called “gravatazal”, “caraguatal” or “gravatá-do-banhado” (in Portuguese) and, due to the close relationship of the Straight-billed Reedhaunter with this ecosystem, its popular scientific name has been recently changed from “junqueiro-do-bico-reto” to “arredio-do-gravatá” (Bencke et al. 2010). The species has no known migratory movements and its geographical distribution is limited to southernmost Brazil –in the Santa Catarina (Fontana et al. 2008) and Rio Grande do Sul states– and in neighboring countries; e.g., Uruguay (Aldabe et al. 2009) and Argentina (Chebez et al 2011). In all these countries, the Straight-billed Reedhaunter is commonly included in Red Lists and the species are globally classified as near-threatened (Aldabe et al. 2009; Chebez et al. 2011; BirdLife International 2012; MMA 2014; Rio Grande do Sul 2014). Very little information on the biometry of nests has been reported for the species (López-Lanús 2016). Furthermore, there no up-to-date reflections exist on the factors that determine the orientation of the opening of this species’ nests and their spatial location within patches. In this work, we present information on the biometry, cardinal orientation and spatial distribution of three Straight-billed Reedhaunter nests in southernmost Brazil, along with comments on this species’ ecology.

## Methods

*Study area and data collection*. Nests were located in the municipality of Bagé, southernmost Brazil (S 31° 17’ 26,4’ and W 54° 09’ 32,7’). This region faithfully represents the Pampa Biome landscape and livestock is the main economic activity there (Pimentel 1940; Rambo 1959; IBGE 2014). In September 2012 we searched nests in the wetlands dominated by Eryngo located on the “Arvorezinha” stream border. Three nests were recorded. For each nest, information about biometrics, location within the patch (border or inland) and the cardinal orientation of chamber opening was collected. Nest measurements were: 1) plant species and height of support plant; 2) distance from the ground to the bottom base nest; 3) vertical and horizontal diameter of the chamber opening; 4) depth egg chamber. After 20 days, nests were reassessed, and information about abandonment and number of eggs and individuals was obtained.

## Results

All nests were located on the edge of patches, on higher and dry sections of the ground (transition between patch and adjacent fields). Two nests were oriented to the north and one was oriented to the east. All nests had two eggs each. In the next month, juveniles in the proximity of two nests, and a nest abandoned with intact two eggs, were observed. Interestingly, the habitat of one of the successfully reproductive nests was totally destroyed in the incubation phase, but the parents remained until juveniles became independent. The mean and standard error (±SE) of the nests measures were: support plant height (108±10.5); distance from ground to bottom of nests (48.1±12.8); vertical outer diameter (17.7±1.5); horizontal outer diameter (12.7±3.7); incubation chamber depth (10.6±2.5); horizontal inner diameter (4.2±0.7) (Table 1).

**Table 1.** Description of the characteristics of the nests observed in our study.

| Characteristic | Nest 1 | Nest 2 | Nest 3 |
| --- | --- | --- | --- |
| Support plant | *E. pandanifolium* | *E. pandanifolium* | *E. pandanifolium* |
| Support plant height (cm) | 100 | 120 | 104 |
| Distance from ground to bottom of nests (cm) | 60 | 50 | 34.5 |
| Vertical outer diameter (cm) | 16 | 19 | 18.2 |
| Horizontal outer diameter (cm) | 16.9 | 11.7 | 9.6 |
| Incubation chamber depth (cm) | 13 | 8 | 10.8 |
| Horizontal inner diameter (cm) | 4.2 | 5 | 3.5 |
| Number of eggs | 2 | 2 | 2 |
| Reproductive success | 2 | Abandoned | 2 |
| Cardinal orientation | North | East | North |
| Location | Border | Border | Border |

## Discussion

### Biometry, number of eggs and reproductive success

Overall the information found for our three study nests is consistent with the limited information available in the literature (Daguerre 1933; Ricci and Ricci 1984; Barbarskas and Fraga 1998; López-Lanús et al. 1999, López-Lanús 2016). Although the breeding season of Straight-billed Reedhaunter has been documented to fall between September and January (Belton 1994), two of the three recorded nests were found early in September (two eggs in each), which indicates that the breeding period began at least in August.

### Spatial distribution of nests

The three nests were located on the edge of patches. These observations are consistent with four other nests observed in southern Brazil (C. B. ANDRETTI & R. A. DIAS pers. com. to GONÇALVES, March 2015). It is known that wide microclimate variations exist between the interior and the edge of humid ecosystems (Raney et al. 2014), and that relative humidity strongly affects development of eggs (Ar and Sedis 2002). Once the breeding period of Straight-billed Reedhaunter begins in winter months, when the interiors of wetlands are usually flooded, the strategy adopted to build nests on the edge of patches might indicate the effect of factors like relative humidity and temperature. It is also possible that the nests located on the edge of patches constitutes a strategy to avoid the attention of predators during nesting as adults use the dense vegetation in the interior of patches to hunt insects.

### Cardinal orientation of nests

Information about the cardinal orientation of nests in Straight-billed Reedhaunters is scarce. To date, only Barbarskas and Fraga (1998) have reported information on this species’ cardinal orientation (northwardly oriented). However, it is known that the phylogenetic proximity between species is also reflected in the structural similarity of nests, especially in Furnariidae (Zyskowsky and Prum 1999; Irestedt et al 2006; Ohlson et al 2013). Some studies conducted with Furnariidae nests have indicated that incubation chamber orientation does not always follow the same pattern, and that the same species may show structural changes according to the habitat conditions of each breeding season (Mezquida 2004; Greeney 2009). Of the three nests observed in our study, two oriented chamber opening to the north and east. Two other nests observed in southern Brazil in December 2005 were oriented to the south and north-northeast (R. A. Dias pers. com. to GONÇALVES, March 2015). It is known that nest orientation is interpreted as a response to microclimate conditions, especially wind direction and solar radiation (Yanes et al. 1996; Mezquida 2004; Burton 2006; Greeney 2009). Overall, the collected information showed that Straight-billed Reedhaunter’s nest orientation tended to lie between north and east. However, more information by comparing habitat structure and chamber orientation in different habitat situations would clarify the orientation pattern of this species’ nests.

The results presented herein extend the scarce knowledge available about the reproductive biology of the Straight-billed Reedhaunter. This species is distributed in the form of metapopulations (patches spatially separated and connected by biological flows), and no information on the mechanisms that affect this species’ abundance and distribution in areas under anthropic influence is available. Thus we finalize our contribution by encouraging research to measure the negative effects of different anthropic activities on its population dynamics, which will provide better information for the conservation of this species.

## Acknowledgments

M.S.S.G and C.B.A received doctoral fellowships from Capes and CNPQ, respectively, while writing this note. The authors thank Rafael A. Dias for the courtesy of the unpublished data. We are grateful to the Ecossis Company for support while undertaking fieldwork.

